# Coordinate-Based fMRI Meta-Analyses of Episodic Memory Encoding and Retrieval in Depression

**DOI:** 10.64898/2026.04.14.718401

**Authors:** Raphaela Schöpfer, Reut Zabag, Florian Wüthrich, Romy Lorenz, Jutta Joormann, Sina Straub, Jessica Peter

## Abstract

**Background:** Depression is a mood disorder frequently associated with episodic memory impairment. However, it remains unclear whether functional brain activity differs between depressed and non-depressed individuals during encoding or retrieval of autobiographical or non-autobiographical memories. Clarifying these differences is important for refining theoretical models of memory impairment in depression and, potentially, for developing targeted interventions.

**Methods:** We conducted three coordinate-based meta-analyses examining encoding and retrieval of autobiographical and non-autobiographical memory in control participants and individuals with current, remitted, or subthreshold depression, or those at risk for depression. Studies were identified via database searches and analysed using Seed-based d Mapping.

**Results:** We included coordinates from 22 fMRI studies. During encoding, depression was associated with reduced activity in the caudate, the salience network, the frontoparietal executive control network, and motor-related areas (11 studies, N = 542). During non-autobiographical retrieval, depression was associated with higher activity in the right inferior frontal gyrus (7 studies, N = 345). During autobiographical retrieval (10 studies, N = 423), depression was associated with reduced activity in the right insula and fusiform gyrus, alongside increased activity in the left anterior cingulate cortex and the left middle frontal gyrus. Between-study heterogeneity was low and no evidence for publication bias was found.

**Discussion:** Our results indicate that depression may be associated with impaired salience integration during encoding and autobiographical retrieval. In contrast, during non-autobiographical retrieval, increased frontal activity suggests a more vigilant or self-monitoring retrieval mode. Functional brain activity changes in depression therefore appear stage- and content-specific.

## Introduction

In addition to negative affect or loss of interest in nearly all activities, depression is characterized by another important component - cognitive symptoms. Episodic memory dysfunction is common [1,2], affecting both autobiographical memory and memory for experimentally encoded, non-autobiographical information (e.g., assessed in laboratory tasks). Both types of episodic memory rely on successful encoding, during which information is selected, integrated, and bound into a durable memory trace [3]. As retrieval depends on the integrity of these initial representations, disturbances at the encoding stage may compromise later retrieval. Hence, encoding may constitute an upstream stage at which memory-related alterations arise that contribute to downstream retrieval impairments. Understanding encoding processes may therefore be critical for elucidating the mechanisms underlying episodic memory dysfunction in depression.

Behaviourally, however, the quality of encoding can only be inferred from retrieval outcomes. For instance, patients with depression often retrieve fewer items than non-depressed participants in non-autobiographical tasks, such as word lists, and tend to retrieve overgeneralized and less detailed autobiographical memories [4,5]. While such findings indicate episodic memory impairment in depression, they do not directly reveal how information is processed during encoding and retrieval and neither whether the encoding stage or rather the retrieval stage is altered. Investigating the activity patterns underlying these stages using fMRI may provide a more direct characterization of memory-related alterations in depression.

Successful encoding relies on coordinated brain activity in the prefrontal cortex, the medial temporal lobe as well as associated posterior cortical regions [6]. Retrieval - autobiographical and non-autobiographical – engages a largely left-lateralized fronto-parietal network with additional involvement of the temporal cortex and the hippocampal formation [7,8]. Encoding and retrieval activity therefore appear to converge in overlapping regions, consistent with theories positing that successful retrieval depends on reactivation of cortical patterns engaged during encoding [9,10]. If retrieval depends on reinstatement of encoding-related representations, alterations observed during retrieval may partly reflect abnormalities already present during encoding. Consistent with this view, several studies reported that similar brain regions show altered activity in individuals with depression during encoding [11–21], autobiographical retrieval [22–31], and non-autobiographical retrieval [13–16,20,21]. However, these studies have largely examined encoding, autobiographical retrieval, or non-autobiographical retrieval in isolation. Hence, whether these alterations are stage-specific (encoding vs. retrieval) or content-specific (autobiographical vs. non-autobiographical) has not been systematically examined. A very recent meta-analysis investigated autobiographical memory retrieval in depression [32] but did not consider stage- or content-specific differences either. Clarifying these distinctions is important to develop a mechanistic understanding of depression-related memory dysfunction and to identify potential therapeutic targets.

This meta-analysis therefore aimed to characterize functional activity alterations associated with episodic memory in depression across three memory processes: encoding, autobiographical retrieval, and non-autobiographical retrieval. Specifically, we aimed to examine whether depression-related alterations are shared across memory stages and retrieval contents or whether they are specific to particular processes. Comparing the functional brain activity across encoding and both autobiographical and non-autobiographical retrieval allows a process-oriented examination of whether depression-related alterations emerge early during memory formation, persist during content-general retrieval, or become amplified with self-referential content. Such a process-oriented approach may provide important insights into the neurocognitive mechanisms underlying memory dysfunction in depression. Given the limited number of studies directly comparing these memory processes in depression-related populations, our analyses were partly exploratory and intended to generate hypotheses regarding stage- and type-specific functional alterations. In addition, age and sex were examined as exploratory moderators because both factors have been associated with variation in memory-related functional brain activity. Importantly, memory impairment in depression is not restricted to individuals meeting full diagnostic criteria for Major Depressive Disorder. Such impairment has been observed in individuals with subclinical depressive symptoms, in those at risk for developing depression, and in patients with remitted depression [4,33–37]. These findings suggest that memory dysfunction may reflect a broader vulnerability associated with depression, rather than merely a consequence of current depressive symptoms. We therefore included studies in patients with current or remitted depression as well as in individuals with subclinical depressive symptoms or those at risk of developing depression due to familial risk (i.e., across the depression continuum) to capture alterations associated with depression across different stages of vulnerability.

## Methods

### Search Strategy

We conducted three coordinate-based meta-analyses in accordance with the PRISMA guidelines [38]. Two pre-registered analyses (OSF: https://doi.org/10.17605/OSF.IO/T54Y2; https://doi.org/10.17605/OSF.IO/U24DS) examined group differences in memory encoding and autobiographical retrieval across the depression continuum. A third, non-pre-registered and exploratory analysis examined whether depression-related changes in brain activity during retrieval are specific to autobiographical memory or extend to non-autobiographical memory. We systematically searched PubMed, PsycINFO, Embase, and Elicit AI (an AI-assisted literature search tool) to identify studies that compared encoding or retrieval processes between individuals on the depression continuum and control participants. The search was originally conducted up to March 2025 and was updated in June 2026 to identify any newly published studies. We included studies that were available in English, with no restriction on the earliest publication date. We used the following search strings:

((((neuronal[Title/Abstract] OR fMRI[Title/Abstract] OR “functional magnetic resonance imaging”[Title/Abstract] OR neuroimaging[Title/Abstract] OR “brain function”[Title/Abstract] OR “neural mechanism”[Title/Abstract] OR hippocampus[Title/Abstract] OR “prefrontal cortex”[Title/Abstract] OR amygdala[Title/Abstract] OR “functional connectivity”[Title/Abstract])) AND (“episodic memory”[Title/Abstract] OR “memory encoding” [Title/Abstract] OR “memory retrieval”[Title/Abstract] OR “recognition memory”[Title/Abstract] OR “autobiographical memory”[Title/Abstract])) AND (depression[Title/Abstract] OR MDD[Title/Abstract] OR “depressive episode” [Title/Abstract] OR “major depressive episode”[Title/Abstract]))

We adapted the search strings for each database if needed. We also screened reference lists of all included studies to identify additional eligible studies.

### Inclusion and exclusion criteria

We included univariate task-based fMRI studies that reported whole-brain activation contrasts related to encoding, non-autobiographical retrieval, or to autobiographical retrieval and that provided peak voxel coordinates in standard stereotactic space (MNI or Talairach). The studies needed to compare control participants with patients with current or remitted depression, individuals with subclinical depressive symptoms, or with individuals at risk for depression. At risk for depression was defined as individuals with no current or past depressive disorder but with increased vulnerability due to familial risk (i.e., having a parent or twin with a history of depression).

We excluded studies if they:

a. used a region-of-interest (ROI) analysis and no whole-brain analysis;
b. examined resting-state fMRI rather than task-based fMRI;
c. focused only on structural MRI;
d. focused on depression with psychotic symptoms or on other psychiatric disorders;
e. reported centre of mass coordinates instead of peak voxel coordinates;
f. did not provide sufficient information (e.g., no *t*- or *z*-values, or no coordinates)

Two independent raters (RS, RZ) checked studies for inclusion using Covidence systematic review software (Veritas Health Innovation, Melbourne, Australia). Any discrepancies were discussed and resolved by consensus. Interrater agreement was moderate during title and abstract screening (Cohen’s *κ* = 0.59; agreement = 80%) and substantial during full-text screening (Cohen’s *κ* = 0.64; agreement = 83%).

We identified 290 studies and finally included 22 studies (Figure 1) comprising N = 1030 participants. Of these 22 studies, 11 examined encoding, 10 autobiographical memory retrieval, and 7 non-autobiographical memory retrieval.

**Figure 1.**
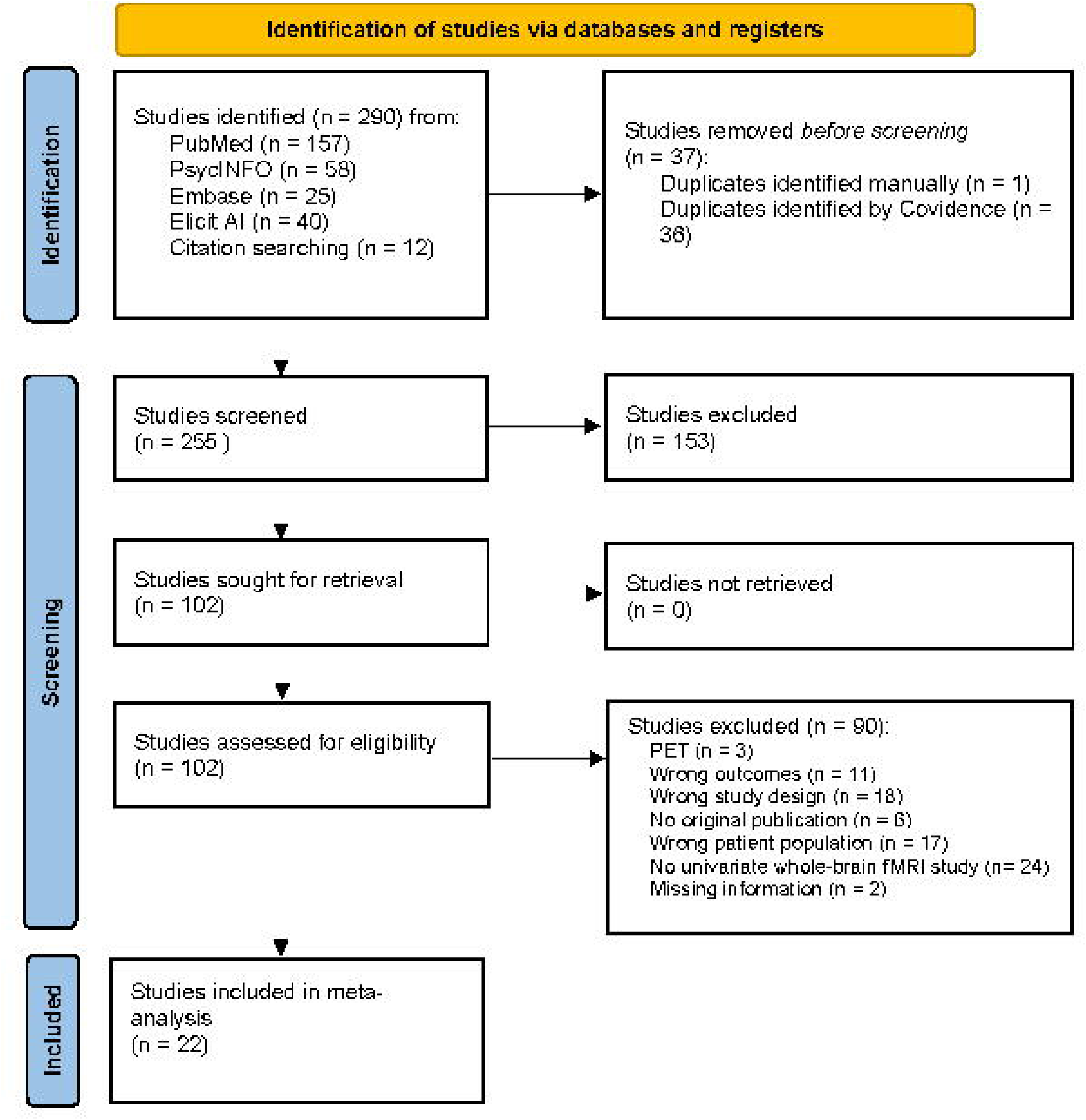
PRISMA flowchart of the study.

### Data Extraction

For each study, two authors (RS, SS) independently extracted MNI or Talairach coordinates, effect sizes (*t*-values), and statistical thresholds. One author (RS) extracted participant information and task descriptions. We contacted authors of studies when information was missing. For one study we received no response, for another study, requested information was unavailable. Hence, these studies had to be excluded. If studies reported *z*-values rather than *t*-values, we converted them using the converter provided on the Seed-based d Mapping website (https://www.sdmproject.com/utilities/?show=Statistics).

Three studies compared three groups (e.g., control participants vs. patients with depression vs. individuals at risk for depression) but did not provide whole-brain, voxel wise pairwise comparisons with extractable peak coordinates or *t*/*z*-maps. If regional summary statistics were available, we combined clinical subgroups into one group using weighted means and pooled standard deviations. We then compared this combined clinical group with control participants using Welch’s *t*-tests to estimate effect sizes. When multiple studies reported results from the same participant sample, only one was included in the meta-analysis to prevent double counting and inflation of effect sizes.

### Seed-Based d Mapping analysis

We performed three coordinate-based meta-analyses using Seed-based d Mapping (SDM) with permutation of subject images (SDM-PSI; version 6.23; www.sdmproject.com) [39]. Preprocessing included the conversion of peak coordinates into standard MNI space and the recreation of whole-brain maps of Hedges’ *g* effect sizes using an anisotropic Gaussian kernel (FWHM = 20 mm, voxel size = 2 mm, anisotropy = 1), restricted to grey matter, following standard SDM procedures for modelling spatial uncertainty of reported peak coordinates [40]. We then calculated mean meta-analytic maps using a random-effects model weighted by study size and variance, and determined statistical significance using permutation tests within the SDM-PSI framework. In line with recommendations for SDM meta-analyses, we used the default threshold for statistical significance (voxel-level *p* < .005 uncorrected, two-sided, minimum cluster extent 10 contiguous voxels). While this threshold does not correct for multiple comparisons, it provides an optimal compromise between sensitivity and specificity [41]. We evaluated between-study heterogeneity using Cochran’s Q statistic and its z-transformed value (Q–Z), τ² (between-study variance), H² (ratio of total to within-study variance), and I² (percentage of true heterogeneity). In addition, we used Egger’s statistic to test for publication bias. Heterogeneity and bias estimates were done using SDM-PSI.

In addition, we conducted three exploratory meta-regressions to examine whether age or sex moderated encoding- or retrieval-related brain activation differences between groups. These analyses were performed as we did not restrict studies to a specific age range, and we aimed to rule out potential confounding effects of age or sex on observed activity differences.

## Results

### Encoding of new information

We included 11 studies comprising N = 542 participants (242 control participants and 300 individuals across the depression continuum). The participants were 36.0 ± 15.2 years old, and 63.7% % of them were female. Encoding tasks focused on words, faces, pictures, or on associative learning (Supplementary Table S1).

We found higher activity in control participants compared to individuals across the depression continuum in the right caudate, the bilateral insula, the right precentral gyrus, the left superior frontal gyrus and the left inferior parietal lobe (Table 1, Figure 2A). Effect sizes were small to medium (Hedges’ *g* ranging from -.456 to -.314). Heterogeneity was low (τ^2^ ≤ .022; I^2^ ≤ 16 %) and not significant (Q–Z < 1.96), indicating consistency between studies. We found no evidence to suggest publication bias (Egger’s regression tests, all *p* > .288; Supplementary Table S4). The reverse contrast (i.e., depression > health) did not yield any significant results.

**Table 1.** Significant differences in brain activity between control participants and the depression continuum during encoding of new information.

| Region | MNI coordinate (x,y,z) | Cluster size | SDM-Z | p-value | Direction | Hedges' g |
| --- | --- | --- | --- | --- | --- | --- |
| Right caudate | 12,-6,16 | 137 | -3.794 | <.001 | control > depression | -.456 |
| Right insula | 38,16,-4 | 131 | -3.617 | <.001 | control > depression | -.417 |
| Right precentral gyrus | 30,-6,56 | 101 | -3.729 | <.001 | control > depression | -.432 |
| Left insula | -40,-14,4 | 45 | -3.030 | .001 | control > depression | -.351 |
| Left superior frontal gyrus | -2,46,-6; 0,60,-8 | 20; 10 | -2.831; -2.873 | .002; .002 | control > depression | -.341;—-.335 |
| Left inferior parietal lobe | -30,-58,42 | 15 | -2.869 | .002 | control > depression | -.313 |

*Abbreviations:* MNI = Montreal Neurological Institute, SDM-Z = Seed-based d mapping z value.

We then examined whether age or sex moderated the observed group differences in brain activity and found no significant effect of age. In contrast, sex significantly moderated the observed group differences in right precentral gyrus activity. Studies with a higher proportion of female participants showed greater differences in encoding-related right precentral gyrus activity between both groups (Supplementary Table S5).

### Retrieval of non-autobiographical memories

For non-autobiographical retrieval, we included 7 studies comprising N = 345 participants (161 control participants and 184 individuals across the depression continuum). They were 32.1 ± 11.1 years old and 65.7 % of them were female. Six studies used a recognition task and one study used a cued-recall task (Supplementary Table S2).

We found higher activity in the right inferior frontal gyrus in the depression group compared to control participants (Table 2, Figure 2B). The effect size was small to medium (Hedges’ *g* = .463). Between-study heterogeneity was low (τ^2^ = 0.013; I^2^ = 8.722 %), and not significant (Q–Z < 1.96), indicating consistency between studies. We found no evidence to suggest publication bias (i.e., Egger’s *p* = .963; Supplementary Table S6). The reverse contrast (i.e., health > depression) did not lead to any significant results.

**Table 2.**
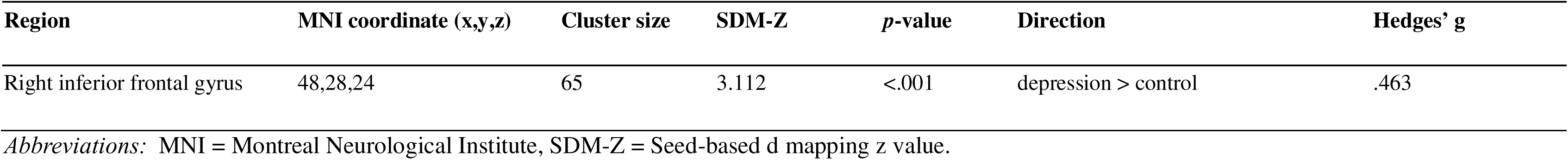
Significant differences in brain activity between control participants and the depression continuum during retrieval of non-autobiographical memories.

We then examined whether age or sex moderated the observed group difference in brain activity. We found no significant moderating effect of age or sex.

### Retrieval of autobiographical memories

For autobiographical retrieval, we included 10 studies comprising N = 423 participants (199 control participants and 224 individuals across the depression continuum). They were 31.3 ± 7.6 years old and 67.0 % of them were female. The studies used different paradigms, including cue-word recall, autobiographical memory recognition, script listening, and photo-based recall (Supplementary Table S3). We found higher activity in the left anterior cingulate and in the left middle frontal gyrus in the depression continuum compared to control participants. There was also lower activity in the right insula and the right fusiform gyrus in the depression group compared to control participants (Table 3; Figure 2C). Effect sizes were small to medium (Hedges’ *g* ranging from -.405 to .539). Between-study heterogeneity was low (τ^2^ ≤ 0.030; I^2^ ≤ 18%) and not significant (all Q–Z < 1.96). We found no evidence to suggest publication bias (Egger’s regression test, all *p* > .229; Supplementary Table S7).

**Table 3.**
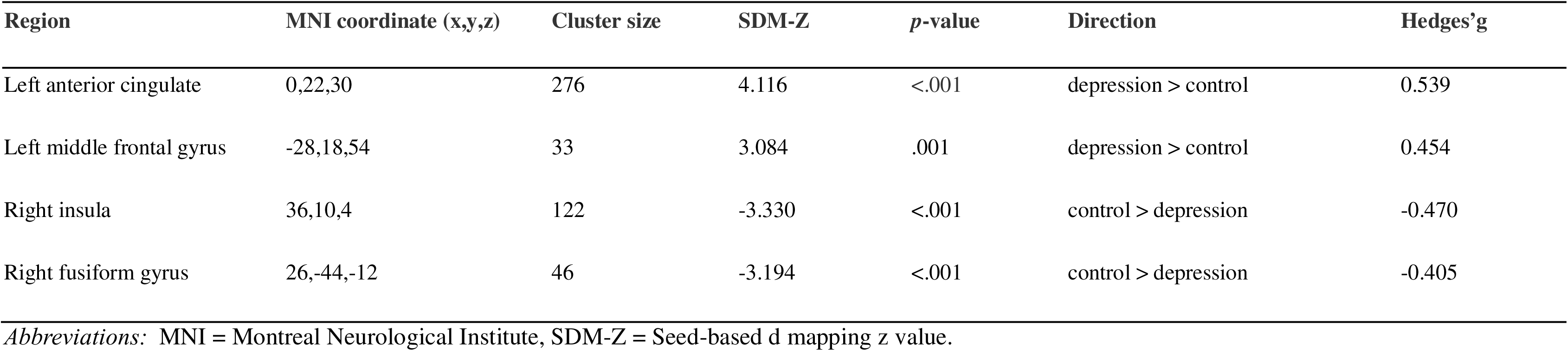
Significant differences in brain activity between control participants and the depression continuum during autobiographical retrieval.

| Region | MNI coordinate (x,y,z) | Cluster size | SDM-Z | p-value | Direction | Hedges'g |
| --- | --- | --- | --- | --- | --- | --- |
| Left anterior cingulate | 0,22,30 | 276 | 4.116 | <.001 | depression > control | 0.539 |
| Left middle frontal gyrus | -28,18,54 | 33 | 3.084 | .001 | depression > control | 0.454 |
| Right insula | 36,10,4 | 122 | -3.330 | <.001 | control > depression | -0.470 |
| Right fusiform gyrus | 26,-44,-12 | 46 | -3.194 | <.001 | control > depression | -0.405 |

*Abbreviations:* MNI = Montreal Neurological Institute, SDM-Z = Seed-based d mapping z value.

We then examined whether age or sex moderated the observed group differences in brain activity. Age significantly moderated the group difference in left superior frontal lobe activity. Higher age was associated with a reduction in the difference in activity between control participants and individuals across the depression continuum. In other words, group differences became smaller with increasing age (Supplementary Table S8). Sex significantly moderated group differences in activity in the left middle frontal gyrus. Studies with a higher proportion of female participants showed greater differences in recall-related left middle frontal gyrus activity between both groups (Supplementary Table S9).

### Conjunction map of all memory processes

As an exploratory analysis, we then created a conjunction map to identify differences in brain activity that are shared between memory stages (i.e., overlaying thresholded maps from all meta-analyses; voxel-level *p* < .005, uncorrected; cluster extent ≥ 10 voxels). We found no significant voxel overlap. However, clusters located within the right insula were in close proximity, indicating consistent hypoactivation across the depression continuum during both encoding and autobiographical memory retrieval (Figure 2D).

**Figure 2.**
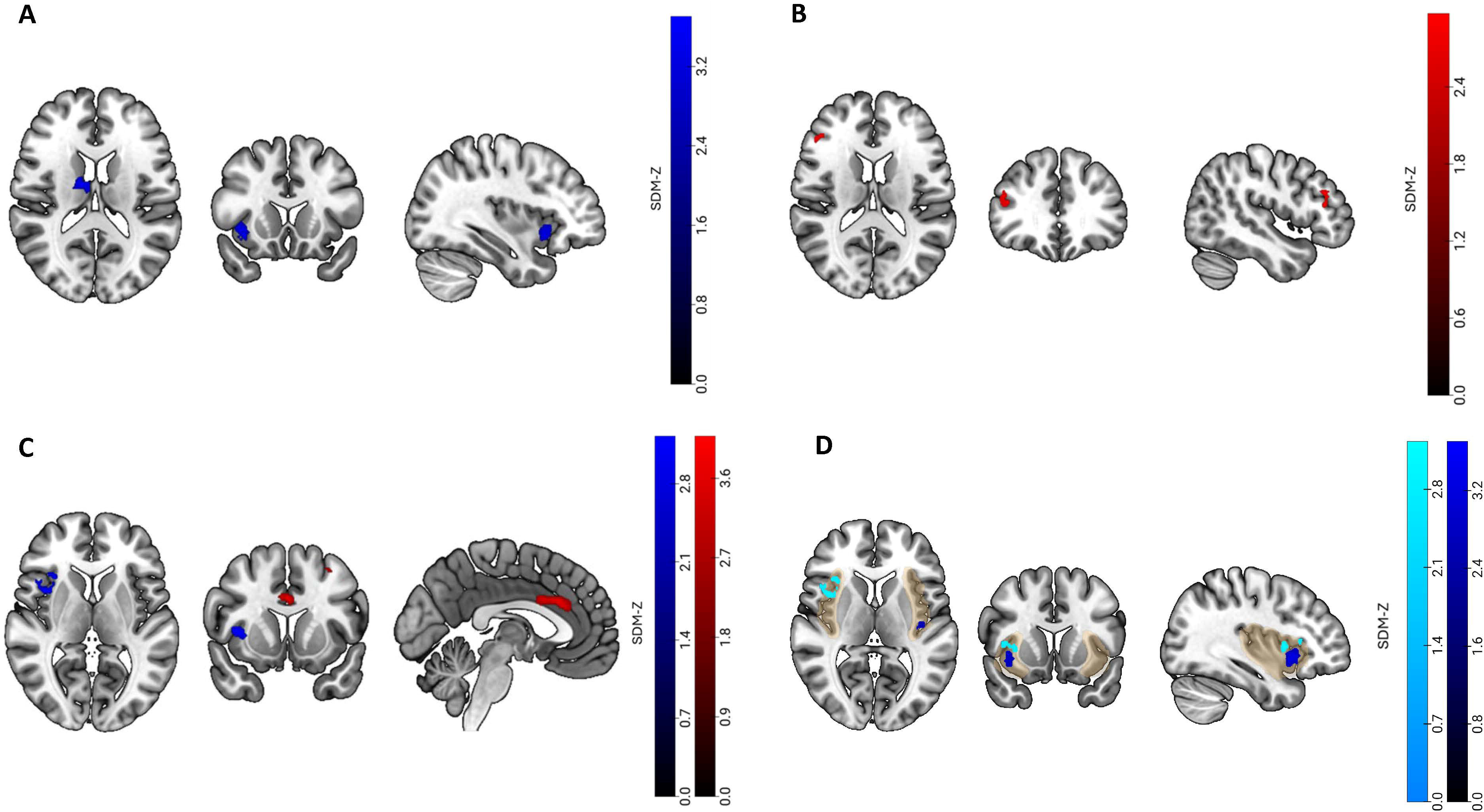
Functional brain activity differences in individuals across the depression continuum compared to control participants during (A) encoding of new information, (B) non-autobiographical memory retrieval, or (C) autobiographical memory retrieval. (D) Adjacent clusters in spatial proximity in the right insula during encoding (dark blue) and autobiographical retrieval (light blue). The statistical threshold was set at *p* < .005, uncorrected, with a minimum cluster extent of 10 voxels. Colour bars represent Seed-based d Mapping z-values. Red denotes increased activity in depression, blue reduced activity.

## Discussion

The goal of the present study was to characterize changes in functional brain activity across the depression continuum during episodic memory, and to explore how these changes vary by memory stage and type. Successful retrieval depends on the quality of initial encoding, during which incoming information is evaluated for relevance, prioritized, and integrated into a coherent representation. Effective encoding relies on executive control that supports elaboration and organization as well as on attention to prioritize information for enhanced processing [42,43].

Across the depression continuum, we observed reduced brain activity during encoding in the right caudate, bilateral insula, left superior frontal gyrus, left inferior parietal lobule, and right precentral gyrus. These regions are key components of the salience network and frontoparietal control network that support the allocation of cognitive resources during encoding. The insula, a core node of the salience network, responds to salient information and supports subsequent memory by flagging important events and by promoting executive resources during encoding [44,45]. In addition, the dorsal part of insula subregions may be important for relevance evaluation of a stimulus to then directly influence hippocampal activity [46] as proposed by a recent intracranial EEG study. Lower activity in the insula across the depression continuum may therefore reflect a reduced capacity to detect and prioritize relevant information, limiting the engagement of integrative processes necessary for forming strong memory traces. Reduced activity in frontoparietal regions may further reflect impaired attentional regulation. reducing the ability to focus on relevant information. Alternatively, this pattern may reflect reduced task engagement or effort, as several identified regions (e.g., the insula and frontoparietal regions) overlap with the multiple-demand network, a domain-general system supporting cognitive control and responses to task demands [47]. Lower activity in the right caudate may reflect reduced goal-relevant processing, including the selection and organization of task-relevant information [48,49]. Together, these findings may indicate that depression is associated with reduced salience processing and executive engagement already at the earliest stage of memory processing, potentially leading to weaker memory traces and less detailed and emotionally salient representations.

During autobiographical retrieval, reduced activity in the right insula persisted across the depression continuum and extended to the right fusiform gyrus. During retrieval, the insula may contribute to the evaluation of personally meaningful experiences by processing interoceptive signals [8,44,45]. The fusiform gyrus may support the reactivation of details that underlie a vivid recollection of personal events [50]. Lower activity in these regions during autobiographical retrieval may therefore compromise a detailed, specific memory reactivation. This, in turn, may lead to overgeneralized autobiographical memories that are less vivid and personally salient. Besides reduced activity in the insula and the fusiform gyrus, we found increased activity in the left anterior cingulate cortex and the left middle frontal gyrus across the depression continuum, pointing to a functional imbalance within the salience network of which the anterior cingulate cortex and the insula are core nodes. This imbalance may alter the coordination of network nodes or may lead to unstable network dynamics. Examining whether this interpretation is supported will require studies that directly assess large-scale network organization and connectivity. Since autobiographical memory retrieval mostly involved a silent free recall, an increase in brain activity across the depression continuum may also indicate increased regulatory demands during the reconstruction of autobiographical memories. Free recall, even when silent, requires a strategic reconstruction of memories when there are no cues that prompt retrieval. Individuals across the depression continuum may need more prefrontal control and monitoring to retrieve even when memories are not very specific (i.e., overgeneralised). Another explanation may be that greater top-down control is needed to reconstruct memories while managing competing negative thoughts.

In contrast to autobiographical retrieval, non-autobiographical retrieval was not associated with reduced activity in the insula or fusiform gyrus but rather with increased activity in the right inferior frontal gyrus, a brain region related to executive control, monitoring, and response selection. Similar to autobiographical retrieval, higher activity in frontal brain regions may indicate increased regulatory demands during the reconstruction of memories. However, non-autobiographical retrieval was largely recognition-based and therefore less demanding than free recall as cues prompt retrieval. In addition, recognition tasks usually lead to *weaker* functional brain activity than, say, free recall tasks. Hence, task difficulty or increased regulatory demands may not be the only reason for an increase in brain activity. Instead, stronger inhibition or monitoring needs, for example, to suppress intrusive thoughts, may play an important role. Hence, depression may alter control dynamics even when memory demands are low [51].

Together, our study highlights the salience network – particularly the insula - as a hub in depression, consistent with previous reports of structural and functional dysfunctions [52–59] or even network reorganisation [60]. An imbalance in the salience network, and in the insula in particular, may contribute to less efficient switching between the default mode network and the executive control network, impairing the brain’s ability to mark memories as important during encoding. This may lead to weaker, less distinct memory traces, contributing to overgeneralized autobiographical memories, in line with previous reports in depression [5,33]. Depression may therefore not erase memories but instead reduce their emotional specificity and vividness. Importantly, during autobiographical and non-autobiographical retrieval, regions supporting cognitive control were more active across the depression continuum. This may reflect greater effort to manage interference from negative thoughts to retrieve memory details despite a known tendency towards overgeneralisation.

Our study has some limitations. First, the number of eligible studies was relatively small, and participants were drawn from across the depression continuum, including individuals with current or remitted depression, those with subthreshold symptoms, or individuals at risk for depression. Due to the limited number of included studies, we were not able to perform subgroup analyses to examine whether activity changes were specific to patients with current depression. Second, the included studies used similar but not identical tasks to assess episodic memory. For the evaluation of non-autobiographical retrieval recognition tasks were mostly used, while autobiographical retrieval included free recall, cue-word recall, and recognition tasks. Despite these methodological differences, similar patterns of increased brain activity were observed across retrieval conditions, indicating that the findings are not solely driven by task-specific demands. In addition, several autobiographical retrieval studies were done by the same research group and used similar paradigms, which may have reduced methodological diversity. However, we did not find evidence of substantial between-study heterogeneity. Third, not all studies matched groups on education, IQ, or general cognitive abilities, and task engagement was not consistently controlled. Fourth, medication may have influenced our results. We extracted medication status from all included studies (Supplementary Table S10), which ranged from unmedicated samples to medicated patient groups. Although medication effects cannot be ruled out, the convergence of our findings across studies with varying medication profiles indicate that medication alone is unlikely to account for the observed patterns. Fifth, our analyses were based on group-level coordinate data. Hence, we could not determine the precise large-scale network affiliation of each identified region at the individual level. Accordingly, interpretations regarding the involvement of specific large-scale networks should be considered tentative and require confirmation using connectivity-based approaches.

## Conclusion

Our study revealed that depression-related functional activity changes vary across memory encoding and retrieval but share convergent insula hypoactivation pointing to disrupted salience processing as a potential mechanism underlying episodic memory dysfunction in depression. These findings may point to potential targets for interventions aimed at improving memory functions. For encoding-related changes in the salience network or frontoparietal regions, attention or memory training may be beneficial. The insula is a promising target for addressing memory dysfunction, for example by using non-invasive brain stimulation. Although direct modulation of the insula remains challenging, interventions that enhance its functional connectivity with other regions, such as the right superior temporal gyrus [61], may hold promise. Another potential approach is real-time fMRI neurofeedback designed to increase insula engagement during episodic memory formation. Future studies may evaluate the efficacy and feasibility of these interventions. In addition, precision functional mapping may characterize network alterations in individuals across the depression continuum and guide targeted interventions.

## Data Availability

The metadata underlying this meta-analysis, including peak coordinates and associated t-values extracted from all included studies, are openly available on the Open Science Framework (OSF) repository at: https://osf.io/wyx76/overview and https://osf.io/mqj3y/overview

## Declaration of generative AI and AI-assisted technologies in the manuscript preparation process

During the preparation of this work the authors used ChatGPT in order to improve grammar and to assist with language editing as well as Elicit AI in order to help with literature search. After using these tools, the authors reviewed and edited the content as needed and take full responsibility for the content of the published article.

