## Supplementary Material for "Coordinate-Based fMRI Meta-Analyses of Episodic Memory Encoding and Retrieval in Depression"

**Table S1.** Studies included in the meta-analysis comparing encoding-related brain activity across the depression continuum and control participants.

| **Study** | **N (control)** | **N (depression)** | **Task** | **Contrast** |
| --- | --- | --- | --- | --- |
| Arnold et al (2011) | 14 | 14 | Encoding of emotional words | Hits > misses |
| Dietsche et al. (2014) | 23 | 23 | Encoding of neutral faces and scrambled pictures | Neutral faces > scrambled pictures |
| [Güleç](https://pubmed.ncbi.nlm.nih.gov/?term=G%C3%BCle%C3%A7+ZN&cauthor_id=41383893) et al (2025) | 13 | 15 | Associative learning (face-name) | Face–name encoding > baseline |
| Holt et al. (2016) | 30 | 56 | Self-referential word memory task | Hits > misses |
| Kassel et al. (2016) | 40 | 42 | Semantic list learning | Encoding > silent rehearsal |
| Roberson-Nay et al. (2006) | 10 | 23 | Face encoding | Successful encoding > unsuccessful encoding |
| Van Eijndhoven et al. (2011) 20 20 Source memory recollection pictures Recollection attempt (Item only > misses) | 20 | 40 | Source memory recollection (pictures) | Source hits > Item hits |
| Van Wingen et al. (2010) | 18 | 28 | Encoding of happy or sad faces | Hits > misses |
| Weisenbach et al. (2014) | 23 | 24 | Semantic list learning | Encoding > silent rehearsal |
| Werner et al. (2009) | 11 | 11 | Associative learning (face-profession) | Encoding > viewing head template |
| Wolfensberger et al. (2008) | 40 | 24 | Encoding and judgement of emotional words | Encoding > attention task |

*Abbreviation: N = Number.*

**Table S2.** Studies included in the meta-analysis comparing non-autobiographical retrieval in control participants and individuals across the depression continuum.

| **Study** | **N (control)** | **N (depression)** | **Task** | **Contrast** |
| --- | --- | --- | --- | --- |
| Dietsche et al. (2014) | 23 | 23 | Recognition of faces | Recognition > scrambled-images |
| Güleç et al (2025) | 13 | 15 | Recognition of face and name | Recognition > Baseline |
| Holt et al. (2016) | 30 | 56 | Recognition of words | Hits > correct rejections |
| Miskowiak et al. (2018) | 26 | 27 | Recognition of words | Hits > correct rejections |
| Van Wingen et al. (2010) | 18 | 28 | Recognition of faces | Hits > misses |
| Werner et al. (2009) | 11 | 11 | Associative retrieval (face-profession) | Retrieval > non-memory perception task |
| Wolfensberger et al. (2008) | 40 | 24 | Recognition of words | Recognition > attention task |

*Abbreviation: N = Number.*

**Table S3.** Studies included in the meta-analysis comparing autobiographical memory retrieval between control participants and individuals across the depression continuum.

| **Study** | **N (control)** | **N (depression)** | **Task** | **Contrast** |
| --- | --- | --- | --- | --- |
| Gillard et al. (2023) | 18 | 18 | Listening to pre-recorded autobiographical memories followed by silent imagery to re-experience them | Positive/negative autobiographical memories > neutral memories |
| Joormann et al. (2012) | 27 | 20 | Silent recall of self-generated positive autobiographical memories | Positive autobiographical memories > fixation cross |
| Parlar et al. (2018) | 15 | 15 | Silent recall of pre-scan-generated autobiographical events in a sequence, cued by on-screen titles | Positive autobiographical memories > odd number detection |
| Whalley et al. (2012) | 15 | 15 | Recognition of words/short sentences extracted from previously written first-person narratives or from unrelated narratives | Own memories > other memories |
| Wu et al. (2020) | 18 | 16 | Viewing of old public photos followed by button press to indicate whether the photo evoked an autobiographical memory (yes/no), followed by rating of the degree of recall | High recall > low recall |
| Young et al. (2012) | 14 | 12 | Viewing of positive, negative, and neutral cue words, followed by button press when an autobiographical memory was recalled | Autobiographical memory > subtraction task |
| Young et al. (2017) | 40 | 40 | Silent recall of autobiographical memories in response to positive, negative, and neutral cue words | Autobiographical memory > example generation task |
| Young et al. (2016) | 16 | 16 | Silent recall of autobiographical memories in response to positive, negative, and neutral cue words | Autobiographical memory > example generation task |
| Young et al. (2014) | 16 | 32 | Silent recall of autobiographical memories in response to positive, negative, and neutral cue words | Autobiographical memory > example generation task |
| Young et al. (2015) | 20 | 40 | Silent recall of autobiographical memories in response to positive, negative, and neutral cue words | Autobiographical memory > example generation task |

*Abbreviation: N = Number.*

**Table S4. Heterogeneity and publication bias statistics for significant clusters in the meta-analysis on memory encoding.**

| **Region** | **τ²** | **H²** | **I² (%)** | **Q²** | **Q-Z** | **Egger’s p** |
| --- | --- | --- | --- | --- | --- | --- |
| Right caudate | 0.019 | 1.163 | 14.047 | 4.419 | -1.706 | 0.870 |
| Right insula | 0.001 | 1.088 | 8.064 | 6.795 | -0.897 | 0.912 |
| Right precentral gyrus | 0.011 | 1.103 | 9.309 | 5.007 | -1.482 | 0.914 |
| Left insula | 0.012 | 1.109 | 9.825 | 7.676 | -0.649 | 0.346 |
| Left superior frontal gyrus | 0.022; 0.017 | 1.202; 1.163 | 16.804; 13.990 | 8.879; 7.631 | -0.340; 0.662 | 0.288; 0.418 |
| Left inferior parietal lobule | 0.000 | 1.003 | 0.333 | 4.807 | -1.556 | 0.835 |

*Note*: τ² = between-study variance; H² = ratio of total to within-study variance; I² = percentage of variability due to heterogeneity; Q–Z = standardized Cochran’s Q statistic; Egger’s p = p-value of Egger’s test for small-study effects.

**Table S5. Sex-related effects in the memory encoding meta-analysis.**

|  | **Region** | **MNI coordinate** | **Cluster size** | **SDM-Z** | **p-value** | **Direction** |
| --- | --- | --- | --- | --- | --- | --- |
|  | Right precentral gyrus | 28,-8,56 | 36 | 3.270 | <.001 | ↑ |

*Note*: SDM-Z = z-values derived from seed-based d mapping. Positive SDM-Z values indicate that studies with a higher proportion of female participants showed greater differences in brain activity.

**Table S6. Heterogeneity and publication bias statistics for significant clusters in the meta-analysis on non-autobiographical retrieval.**

| **Region** | **τ²** | **H²** | **I² (%)** | **Q²** | **Q-Z** | **Egger’s p** |
| --- | --- | --- | --- | --- | --- | --- |
| Right inferior frontal gyrus | 0.013 | 1.096 | 8.722 | 1.200 | -1.993 | 0.963 |

*Note*: τ² = between-study variance; H² = ratio of total to within-study variance; I² = percentage of variability due to heterogeneity; Q–Z = standardized Cochran’s Q statistic; Egger’s p = p-value of Egger’s test for small-study effects.

**Table S7. Heterogeneity and publication bias statistics for significant clusters of the meta-analysis on autobiographical retrieval.**

| ***Region*** | | **τ²** | ***H²*** | **I² (%)** | **Q²** | **Q-Z** | **Egger’s *p*** |
| --- | --- | --- | --- | --- | --- | --- | --- |
|  | Left anterior cingulate | <.001 | 1.020 | 1.985 | 1.390 | -2.861 | 0.948 |
|  | Left middle frontal gyrus | .030 | 1.216 | 17.771 | 3.942 | -1.373 | 0.528 |
|  | Right insula | .030 | 1.226 | 18.439 | 5.584 | -0.775 | 0.229 |
|  | Right fusiform gyrus | <.001 | 1.004 | 0.421 | 4.010 | -1.345 | 0.699 |

*Note*: τ² = between-study variance; H² = ratio of total to within-study variance; I² = percentage of variability due to heterogeneity; Q–Z = standardized Cochran’s Q statistic; Egger’s p = p-value of Egger’s test for small-study effects.

**Table S8. Age-related effects in the autobiographical retrieval meta-analysis.**

|  | **Region** | **MNI coordinate** | **Cluster size** | **SDM-Z** | **p-value** | **Direction** |
| --- | --- | --- | --- | --- | --- | --- |
|  | Left superior frontal gyrus | -24,-10,50 | 27 | -3.369 | <.001 | ↓ |

*Note:* SDM-Z = z-values derived from seed-based d mapping. Negative SDM-Z values indicate that group differences became smaller with increasing age.

**Table S9. Sex-related effects in the autobiographical retrieval meta-analysis.**

|  | **Region** | **MNI coordinate** | **Cluster size** | **SDM-Z** | **p-value** | **Direction** |
| --- | --- | --- | --- | --- | --- | --- |
|  | Left middle frontal gyrus | -26,-8,52 | 18 | 2.905 | .002 | ↑ |

*Note:* SDM-Z = z-values derived from seed-based d mapping. Positive SDM-Z values indicate that studies with a higher proportion of female participants showed greater differences in brain activity.

**Table S10. Medication status in all included studies.**

| **Study** | **Included in meta-analysis on:** | **Medication** |
| --- | --- | --- |
| Arnold et al (2011) | Encoding | All participants were medication-free; 8 of 14 remitted patients had taken antidepressants before |
| Dietsche et al. (2014) | Encoding/Non-autobiographical retrieval | 20/23 medicated (antidepressant monotherapy n=13, antidepressant combined therapy n=3, antidepressant plus antipsychotic n=4); 3 unmedicated; medication type not reported |
| [Güleç](https://pubmed.ncbi.nlm.nih.gov/?term=G%C3%BCle%C3%A7+ZN&cauthor_id=41383893) et al. (2025) | Encoding/Non-autobiographical retrieval | All patients received antidepressants; medication types and dosages not reported |
| Gillard et al. (2023) | Autobiographical retrieval | 10/18 medicated (fluoxetine n=3, citalopram n=2, sertraline n=1, mirtazapine n=1, other n=3); 8/18 unmedicated |
| Holt et al. (2016) | Encoding and non-autobiographical retrieval | Final patient sample (n=56) unmedicated |
| Joormann et al. (2012) | Autobiographical retrieval | All participants were medication-free |
| Kassel et al. (2016) | Encoding | 17/42 patients medicated (40.5%); medication type not reported |
| Miskowiak et al. (2018) | Non-autobiographical retrieval | All participants were medication-free |
| Parlar et al. (2018) | Autobiographical memory retrieval | Most patients were medicated, specifics were not reported |
| Roberson-Nay et al. (2006) | Encoding | All participants were medication-free |
| Van Eijndhoven et al. (2011) | Encoding | Depressed patients medication-naïve (20/20); recovered patients medication-free at scan, but 18/20 previously treated with SSRIs/SNRIs and 1 with paroxetine and amitriptyline |
| Van Wingen et al. (2010) | Encoding/Non-autobiographical retrieval | Currently depressed patients: medication-naïve (n=18). Recovered patients: medication-free at scan; previous SSRI/venlafaxine treatment discontinued ≥2 months before scanning (n=17) |
| Whalley et al. (2012) | Autobiographical memory retrieval | 10/15 currently depressed patients: SSRI/SNRI treatment; 1 with lithium as add-on |
| Weisenbach et al. (2014) | Encoding | 78% of patients on medication; 22% unmedicated. Medicated patients: SSRI/SNRI only (n=7), SSRI/SNRI + other medication (n=7), non-SSRI/SNRI antidepressants (n=3), benzodiazepine only (n=1) |
| Werner et al. (2009) | Encoding/Non-autobiographical retrieval | 73% medicated (6 SSRI, 2 SNRI); lithium augmentation (n=2), benzodiazepines as needed (n=2), antipsychotics (n=3). |
| Wolfensberger et al. (2008) | Encoding/Non-autobiographical retrieval | All participants were medication-free |
| Wu et al. (2020) | Autobiographical memory retrieval | All participants were medication-free |
| Young et al. (2012) | Autobiographical memory retrieval | All participants were medication-free |
| Young et al. (2017) | Autobiographical memory retrieval | All participants were medication-free |
| Young et al. (2016) | Autobiographical memory retrieval | All participants were medication-free |
| Young et al. (2014) | Autobiographical memory retrieval | All participants were medication-free |
| Young et al. (2015) | Autobiographical memory retrieval | All participants were unmedicated at scan; remitted patients were medication-free ≥ 3 months (50% antidepressant-naïve) |

*Abbreviations*: SSRI = selective serotonin reuptake inhibitor, SNRI = selective noradrenaline reuptake inhibitor.
